# Dual role of strigolactone receptor signaling partner in inhibiting substrate hydrolysis

**DOI:** 10.1101/2021.12.01.470725

**Authors:** Briana L. Sobecks, Jiming Chen, Diwakar Shukla

**Affiliations:** Department of Chemical and Biomolecular Engineering, University of Illinois at Urbana-Champaign, Urbana, IL, 61801; Center for Biophysics and Quantitative Biology, University of Illinois at Urbana-Champaign, Urbana, IL, 61801; National Center for Supercomputing Applications, University of Illinois, Urbana, IL, 61801; Beckman Institute for Advanced Science and Technology, University of Illinois at Urbana-Champaign, Urbana, IL, 61801; NIH Center for Macromolecular Modeling and Bioinformatics, University of Illinois at Urbana-Champaign, Urbana, IL, 61801

## Abstract

Plant branch and root growth relies on metabolism of the strigolactone (SL) hormone. The interaction between the SL molecule, *Oryza sativa* DWARF14 (D14) SL receptor, and D3 F-box protein has been shown to play a critical role in SL perception. Previously, it was believed that D3 only interacts with the closed form of D14 to induce downstream signaling, but recent experiments indicate that D3, as well as its C-terminal helix (CTH), can interact with the open form as well to inhibit strigolactone signaling. Two hypotheses for the CTH induced inhibition are that either the CTH affects the conformational ensemble of D14 by stabilizing catalytically inactive states, or the CTH interacts with SLs in a way that prevents them from entering the binding pocket. In this study, we have performed molecular dynamics (MD) simulations to assess the validity of these hypotheses. We used an *apo* system with only D14 and the CTH to test the active site conformational stability and a *holo* system with D14, the CTH, and an SL molecule to test the interaction between the SL and CTH. Our simulations show that the CTH affects both active site conformation and the ability of SLs to move into the binding pocket. In the *apo* system, the CTH allosterically stabilized catalytic residues into their inactive conformation. In the *holo* system, significant interactions between SLs and the CTH hindered the ability of SLs to enter the D14 binding pocket. These two mechanisms account for the observed decrease in SL binding to D14 and subsequent ligand hydrolysis in the presence of the CTH.

## Introduction

Parasitic activity of witchweed, or *Striga*, leads to a $10 billion loss per year of staple food crops and affects the livelihood of 100 million farmers worldwide.^1^ Receptor proteins in *Striga* sense the presence of host crops, such as *Oryza sativa*, via strigolactone (SL) molecules exuded from their roots.^2^ Given that *Striga* germinates at concentrations as low as 2 pM, compared with micromolar concentrations in *At*KAI2, understanding SL behavior in host crops could provide a mechanism for controlling *Striga* proliferation.^3^

SLs are a class of plant hormones involved in branch and lateral root development regulation. ^4–7^ SLs consist of a tricyclic portion (A, B and C rings) and a butenolide ring (D-ring) connected by an enol-ether bridge.^7^ DWARF14 (D14), the SL receptor protein in most angiosperms, is an α-βhydrolase with a Ser-Asp-His catalytic triad and a hydrophobic binding pocket.^8^ When SL enters the binding pocket of D14 in the inactive state, the catalytic serine (S97 in *Oryza sativa* D14) attacks the D-ring, cleaving it from the ABC rings. The catalytic histidine (H247 in OsD14) may covalently bond to the D-ring as the tricyclic structure exits the binding pocket, inducing a closed, or active, conformational state where the lid domain (Helices T1, T2, T3, and T4; see Fig.1) collapses from four helices to three.^9,10^ Activte-state D14 also forms a complex with signaling partners D3 and D53.^10,11^ Following hydrolysis, D14 releases the D-ring component and the D14-D3-D53 complex may undergo degradation, though it is unclear if this degradation occurs after every reaction. ^7^

**Figure 1:**
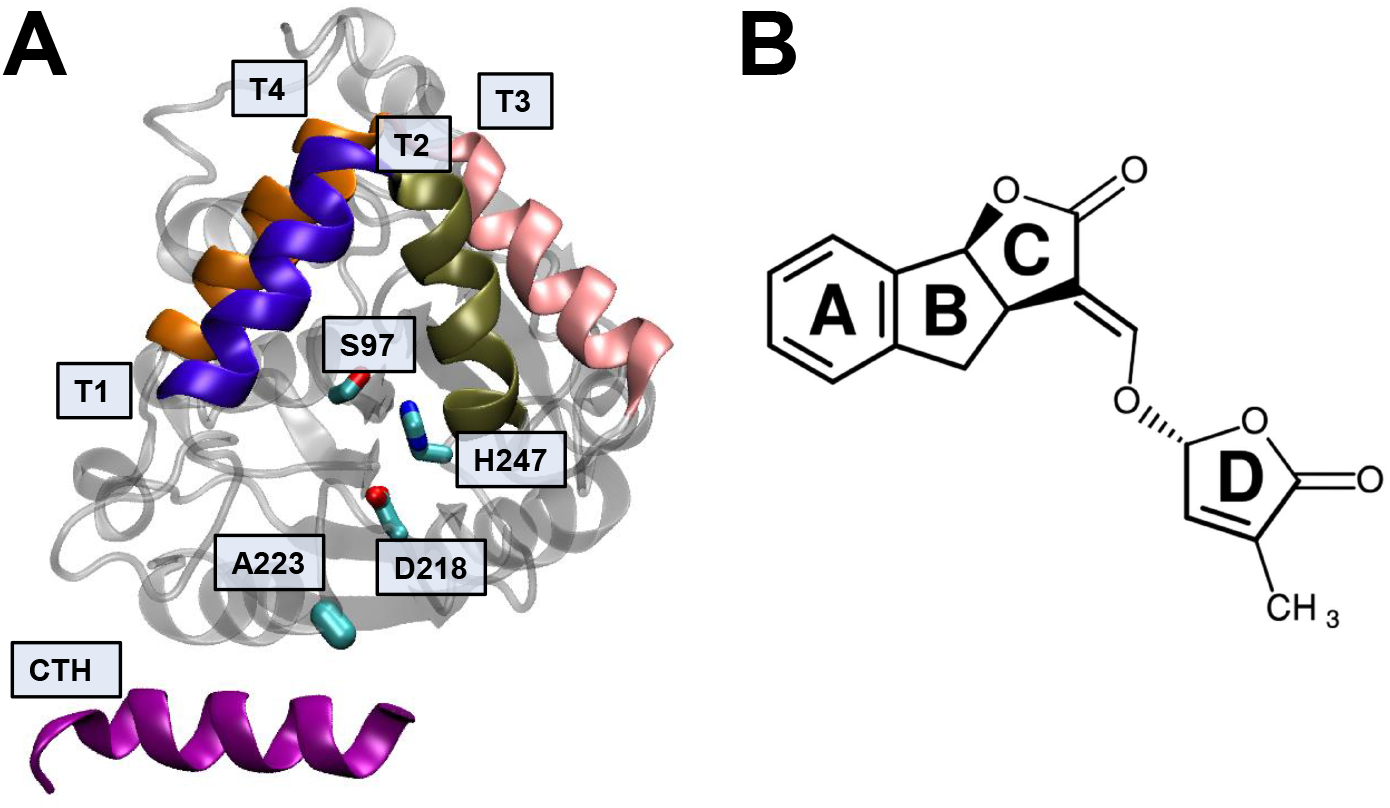
Key structural features of *Os*D14, *Os*D3 C-Terminal Helix (PDB code 6BRT^12^), and strigolactone analogue GR24. (A) The structure of the D14-D3 complex. Major helices on D14 are T1, shown in indigo, T2, shown in tan, T3, shown in pink, and T4, shown in orange. Four important residues are depicted here in cyan. S97, H247, and D218 make up the catalytic triad, which bind GR24, and A223, which binds the D3 CTH in its lowest energy conformation. (B) The structure of GR24, labeled with appropriate ring structures. This is the R-form of GR24 and which is consistent with naturally occuring strigolactones.^7^

SL perception relies on two major signaling partners, an F-box protein (D3 in *O. sativa*) involved in ubiquitination, and a repressor (D53 in *O. sativa*) of SL signaling.^11,13^ A signaling complex forms between D3, D14, and D53, in which D3 association with D14 stabilizes the active, D-ring bound state and decreases the rate of hydrolysis. ^10,11^ Originally, it was believed that D3 and D53 would complex with D14 solely in its activated state; however, recent observations indicate that D3’s C-terminal helix (CTH) can interact with the inactivated form of D14.^7,12^ Furthermore, the presence of the CTH greatly inhibits the rate of D14 SL hydrolysis.^12^ A crystal structure of D14 with the D3 CTH (PDB 6BRT) shows the relative position of the two proteins (Fig.1).^12^ The intact SL molecule may be responsible for activating D14 through a conformational change in the D-loop.^14^ Elucidating the true nature of the D14-D3 interaction may prove key towards understanding the impact of the CTH on SL hydrolysis.

We generated two main hypotheses in regards to the CTH inhibition of strigolactone hydrolysis. One hypothesis is that the presence of the CTH affects the conformational ensemble of the catalytic triad. The CTH is shown in a crystal structure to associate on the outside of the D14 binding pocket such that it does not interact directly with the catalytic triad.^12^ However, CTH interaction with D14 may cause an allosteric effect on triad residue positions or other regions of significance. An allosteric effect from the CTH would make D14 more stable in catalytically inactive conformations, accounting for the observed decrease in SL hydrolysis rate. We tested this hypothesis by running an *apo* molecular dynamics (MD) system with only D14 and the CTH present to evaluate the influence of the CTH on the catalytic triad. The second hypothesis is that the CTH interactions with SLs inhibit the ability of SLs to access the ligand binding site. Given that the CTH does not enter the binding pocket, attraction between SLs and residues on the CTH would prevent the ligand from entering the pocket. This in turn would inhibit SL binding and account for the decreased hydrolysis rate. We tested this CTH-SL interaction with a *holo* MD system containing D14, the CTH, and GR24, a synthetic SL molecule, to evaluate the effect of the CTH on SL binding process.

Using molecular dynamics simulations combined with Markov state models (MSMs) is a popular method to eliminate sampling bias and provide high-resolution, all-atom depictions of protein-ligand interactions beyond static crystal structures.^15–20^ We have recentlt employed this approach for investigating the mechanisms of plant hormone perception and conformational changes in plant proteins.^18–24^ Running a large number of simulations in parallel makes this approach a viable method in terms of efficiency and computing power while still maintaining accuracy of results. ^25–27^ Two previous studies have used MD to analyze α-βhydrolase activity in plant hormone perception. A study examined the binding of the smoke-derived germination stimulant karrikin to its receptor KAI2 in *Arabidopsis thaliana* using steered MD to simulate pulling the ligand away from the binding pocket. ^28^ More recently, our group compared SL perception of D14 versus *Sh*HTL7, the highly-sensitive *Striga* SL receptor, and found several differences in molecular interaction between the two proteins that contribute to the vast discrepancy between concentration sensitivities. ^20^ Here, we used unbiased long-timescale distributed MD systems with approximately 200 μs per system. For simulation construction, we employed adaptive sampling, in which we clustered all simulation frames based on structural features, selected undersampled regions based on a “least-counts” principle, and generated subsequent simulations using frames from the undersampled regions as a starting point. This approach increases efficiency by directing simulations towards unexplored areas while limiting sampling of highly-populated regions.^26,29–31^

In this study, we show that the CTH impacts the rate of hydrolysis by affecting both the active site conformation and SL motion. In the *apo* system, the CTH lowers the free energy barrier between catalytically active and inactive states, creating a higher population within the catalytically inactive state. In the *holo* system, the CTH frequently interacts with GR24, limiting GR24’s ability to enter the binding pocket and bind to the active site. These observations provide insights into the mechanism of SL inhibition by C-terminal helix of D3 that could provide avenues for regulating *Striga* proliferation.

## Results and Discussion

### CTH Decreases the Free Energy Required to Access Catalytically Inactive Conformations

Using our simulation data, we generated free energy landscapes projected onto distance metrics to determine the stability of various interactions within and between the proteins and ligand in these systems. Interactions of the D14 catalytic triad, consisting of residues S97, D218, and H247, determine the likelihood of SL hydrolysis. This is supported by mutagenesis experiments showing that hydrolytic activity of D14 receptors is eliminated when any of the catalytic triad residues are mutated. ^10,14,32^ The *apo* simulations show the lowest free energy conformation is the catalytically active conformation in which the D-loop is attached and the catalytic triad residues are in close proximity (Fig. 2). Two catalytically inactive conformations exist as well, one in which D218 and H247 are separated, and one in which S97 and H247 are separated. The association between the CTH and D14 lowers the free energy barrier between the catalytically active and inactive states by around 2 kcal/mol for the D218-H247 interaction and 3-4 kcal/mol for the S97-H247, creating higher populations in the inactive conformations. Therefore, the CTH contributes to lowered enzymatic activity by altering the conformational ensemble of the catalytic triad.

**Figure 2:**
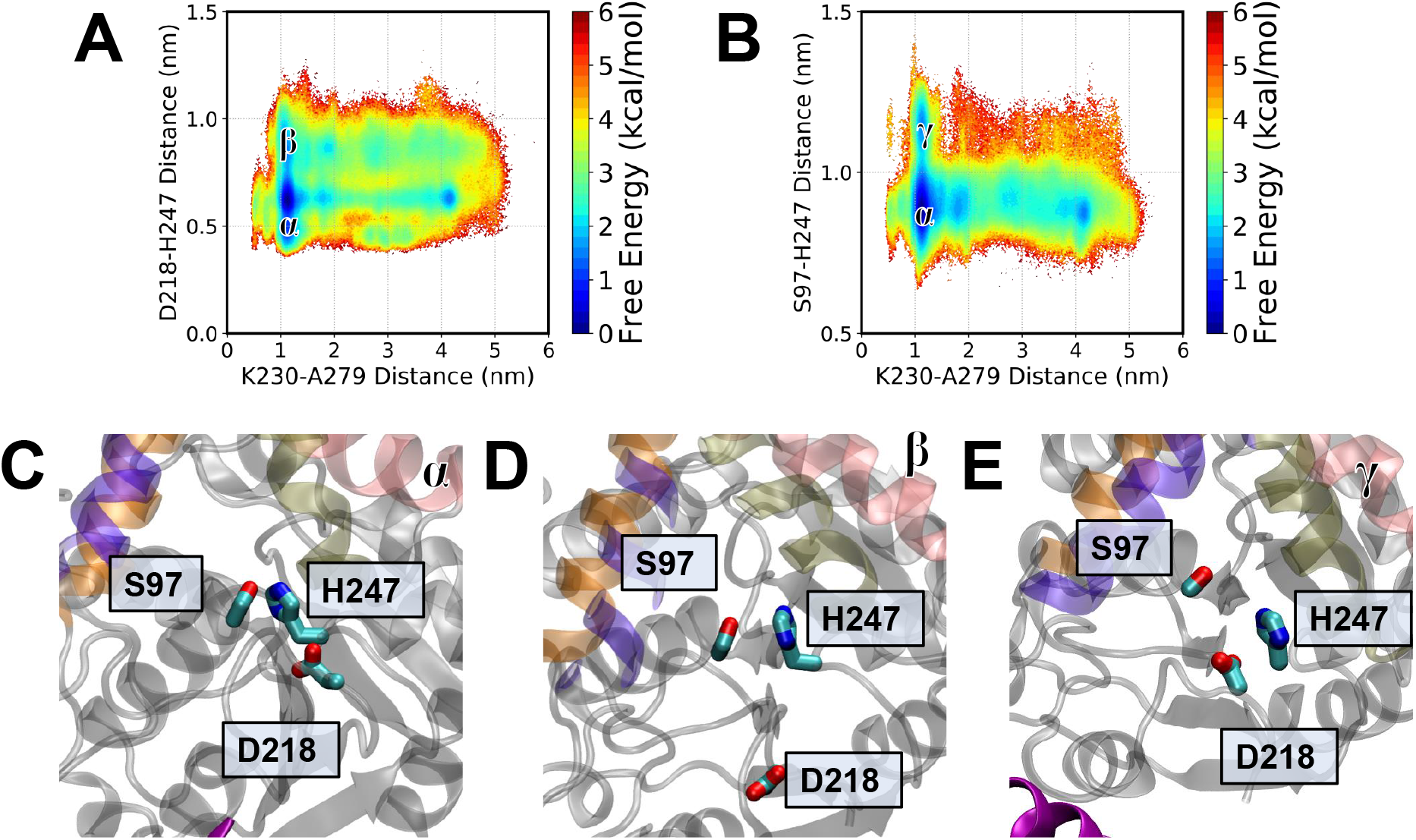
Catalytic triad dynamics versus CTH position. Free energy landscape of CTH dissociation versus the D218-H247 distance (A) and S97-H247 distance (B). (C) Catalytic triad residues in the catalytically active state (α). (D) Catalytic triad conformation in the D-loop detached state, in which D218 and H247 are separated (β). (E) Catalytic triad conformation in which S97 and H247 are separated (γ).

### GR24 Lowers the Free Energy Required for CTH Dissociation

While the CTH allosterically affects the catalytic triad when in the crystal structure position, we also observed the CTH dissociating from this position, which could decrease the stability of the catalytically inactive state. Fig. 3 shows the free energy landscape for interaction between D14 and the CTH in both *apo* and *holo* conditions. The highest stability for both systems was an “associated state” in which the CTH binds to a region of D14 below and to the left of the binding pocket (minima labeled δ). A223 in particular showed a large degree of contact with the CTH. Dissociation from this position requires surmounting an energy barrier of about 3 kcal/mol in *apo* conditions and around 2 kcal/mol in *holo* conditions. In high-energy positions (5 kcal/mol or greater), the CTH is completely dissociated from D14 and floats freely in solution (minima labeled ε). While the increased CTH dissociation from its initial position limits the catalytically inactive state stabilization that occurs from that interaction, this dissociation increases the time in which GR24 interacts with the CTH. A third state of intermediate stability (around 2-3 kcal/mol) exists in which the CTH associates with the lid helices of D14 (minima labeled ζ). This state is around 1 kcal/mol lower in free energy in the *holo* system compared to the *apo* system. In this state, GR24 forms a complex with both lid helices and the CTH, suggesting that the three-way interaction greatly stabilizes this configuration. The association also stabilizes GR24 in a position from which it cannot access the binding pocket. Since GR24 binding inside the pocket is required for hydrolysis, this interaction contributes to the decreased rate of SL hydrolysis in the presence of the CTH. Overall, despite the decrease in allosteric effects, the increased CTH-GR24 interactions decrease SL hydrolysis. When the ligand is present, its interaction with the CTH lowers the free energy barrier for dissociation from D14, which reduces the allosteric stabilization of the catalytically inactive state. However, the presence of the CTH still inhibits hydrolysis due to these GR24-CTH interactions.

**Figure 3:**
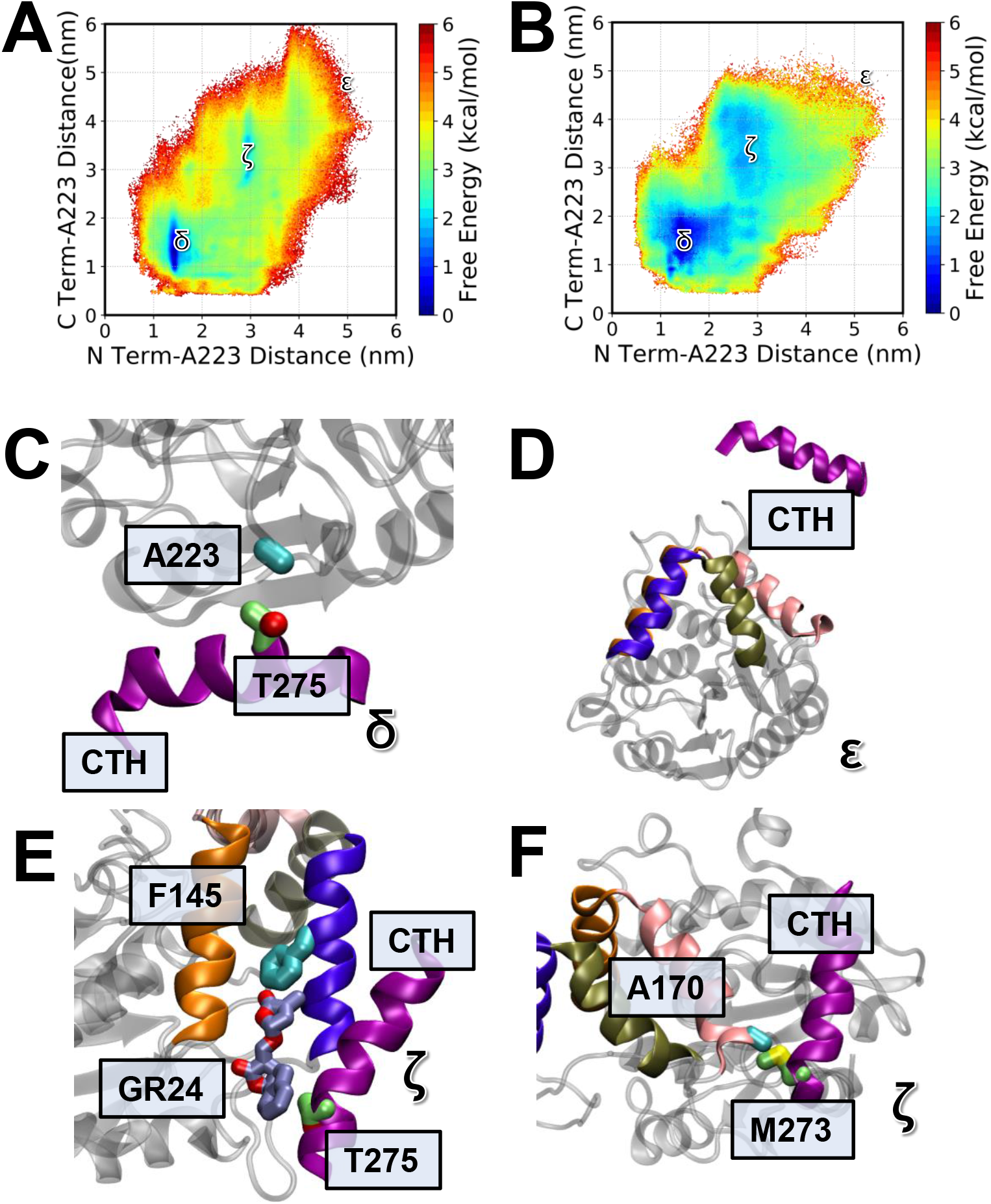
Positions of CTH relative to D14. Free energy landscapes of CTH terminals respective to A223 in *apo* (A) and *holo* (B) systems. (C) CTH and D14 in the associated (δ) state, where CTH residues (lime) are in contact with A223 (cyan) on D14. (D) The CTH (purple) in its high-energy dissociated state (ε). (E) Three-way interaction between GR24 (ice blue), F145 on the T1 helix (cyan) and T275 on the CTH (lime). This cross-interaction stabilizes the complex structure (ζ). (F) Complex formed between D14 T3 helix residue A170 (cyan) and CTH residue M273 (lime). Both lid helix complexes here (ζ) have nearly identical free energy.

### Contact Probability of GR24 with D14 and CTH Residues

While we captured the full binding pathway of GR24 in the presence of the CTH, the ligand favored a few main interactions. Contact probabilities here are defined as the fraction of MSM-weighted total frames in which a given residue is within 4 Å of GR24. The most frequent contacts were with the T1 and T4 helices just to the left of the binding pocket, specifically residues E139, Q143, D146, T193 and K196. These contacts correspond with the three-way D14-D3-GR24 interactions demonstrated in the CTH dissociation analysis. The next major contacts were residues A105 and R108 to the rear of D14. The CTH contained many high-contact residues as well, specifically F274, R278, A279, and L283 (Fig. 4). Furthermore, every residue on the CTH except the C-terminal cap exhibited a contact probability above 4 percent. The total fraction of states in which the ligand came in contact with the CTH was 0.243. The binding probability for GR24 in the presence of the CTH was 0.032, while for our previous simulations on *At*D14 without the CTH, the probability was 0.286.^20^ Using the relation Δ*G* = −*RT* ln *K*, where *K* is the ratio of ligand-bound probabilities, this represents a stability decrease of about 1.31 kcal/mol in the CTH system compared with only D14. The high probability of interaction between the CTH and GR24 indicates a large degree of stability which accounts for the decreased amount of time GR24 spends in the binding pocket for catalysis. Because the CTH never enters the binding pocket, GR24 cannot associate with the CTH and bind to the catalytic triad.

**Figure 4:**
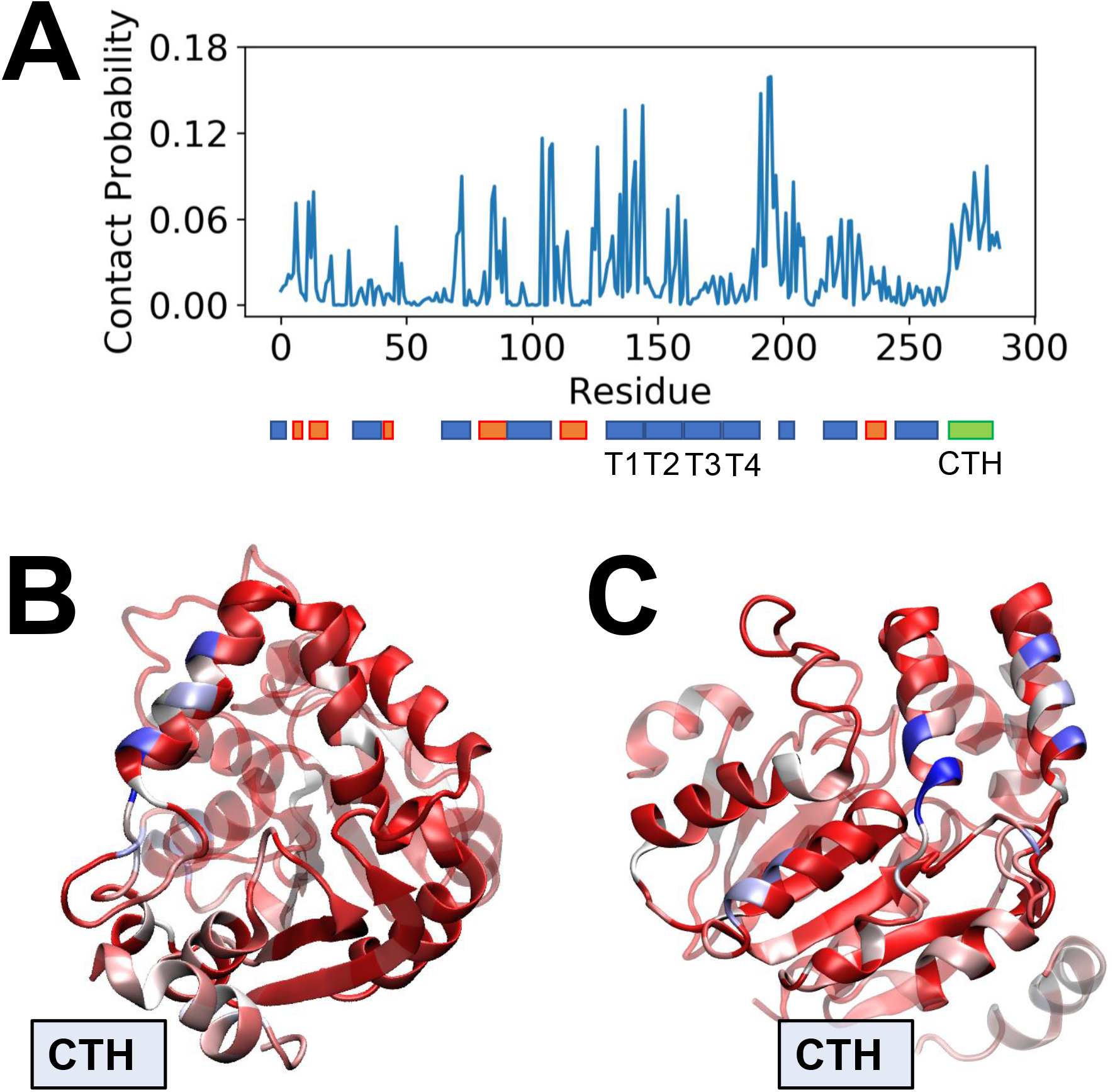
Contact probability of GR24 with protein. (A) Contact probability values for all residues in both D14 and the CTH. Contact probability is the fraction of frames in which the ligand interacts with a given residue. A secondary structure map is given below the graph, with alpha helices in blue, beta sheets in orange, and the CTH in green. (B) Contact probability mapped onto the D14-CTH crystal structure viewed from the front and from the side (C). Highest contact probabilities are colored dark blue, and all probabilities under a 4 percent cutoff are colored dark red, with white representing residues of intermediate probability.

### Transition Path Theory of GR24 Binding

Figure 5 shows the results of transition-path theory (TPT) analysis for the D14-CTH-GR24 system. Mean first passage time (MFPT) was computed between eleven macrostates defined based on the positioning of CTH and GR24 relative to D14. The highest fluxes were into States 5 and 11, which correspond to an unbound GR24 molecule with the CTH either in its associated or dissociated positions, respectively. States in which GR24 anchors to T1/T4 also had significant positive flux. States 4, 8, and 10, corresponding to GR24-CTH interaction, saw a much higher flux than States 1 and 6, in which GR24 was productively bound in the pocket. State 10, with GR24 and CTH associated but dissociated from D14, had one state with a rate constant above 3 μs^−1^ and seven more with a constant above 2 μs^−1^. State 4 (GR24-CTH association with the CTH in its D14-associated state) had eight states with rate constants above 1 μs^−1^. While State 8 (GR24-CTH-T1/T4 in three-way interaction) had fairly low flux, its rate constant values were generally higher than the values for the bound states (State 1 and State 6). The average rate constant value was 0.518 μs^−1^ for State 8, 0.475 μs^−1^ for State 1, and 0.203 μs^−1^ for State 6. Out of all fluxes into the two bound states, only the rate constant from State 5 (unbound GR24, associated CTH) into State 1 was above 1 μs^−1^. Therefore, the fluxes into GR24-CTH associated states were much larger than the fluxes into D14-bound states. This indicates that GR24 spends much more time bound to the CTH than bound to D14. As a result, GR24-CTH interactions are more favorable than GR24-pocket interactions, and more states move here than they do to the bound states. Combined with the high flux into unbound and T1/T4 associated GR24 states, very few states transition into the pocket-bound configurations. This corroborates the decreased rate of GR24 hydrolysis in the presence of the CTH.

**Figure 5:**
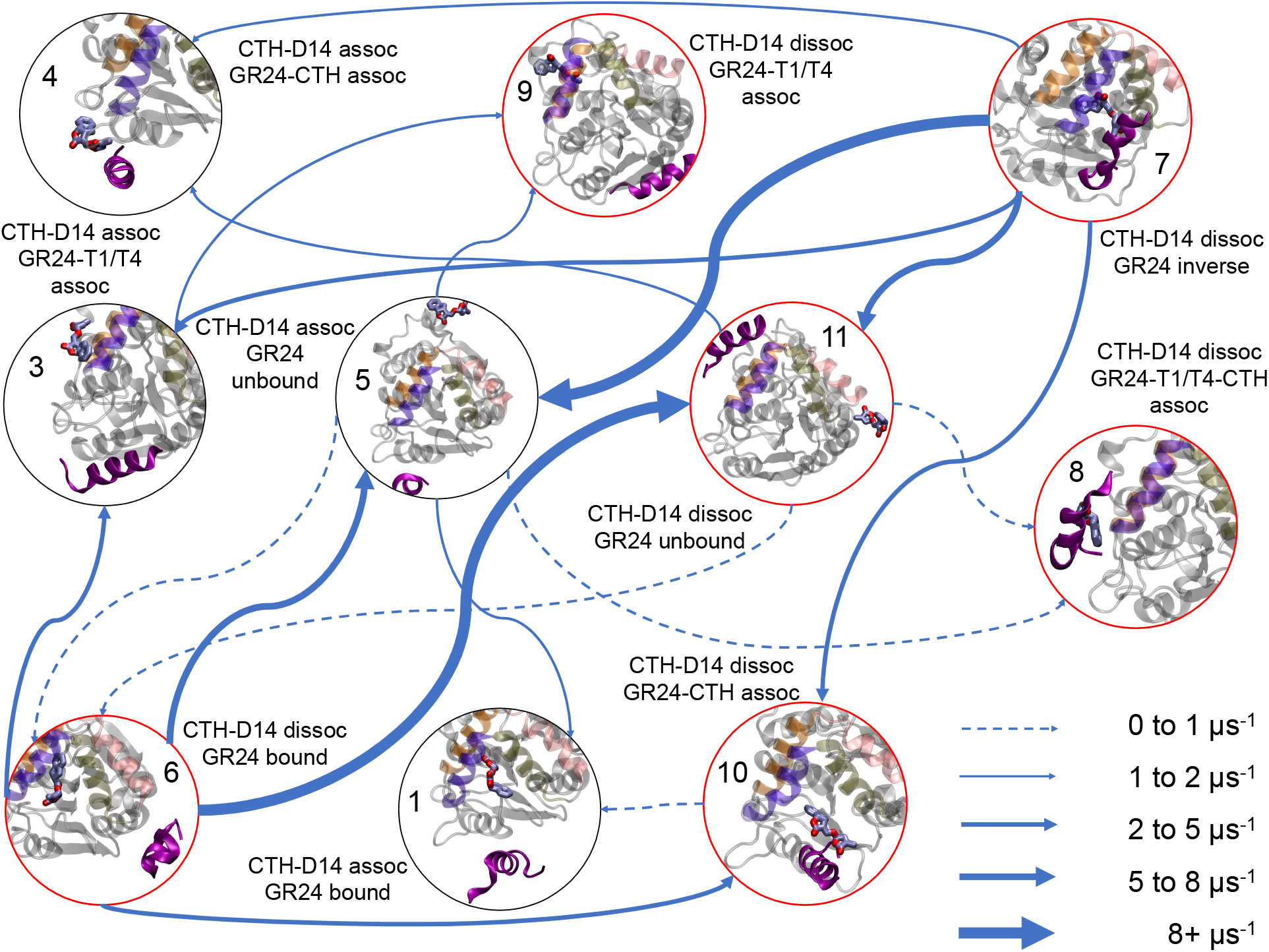
Flux between the major D14-CTH-GR24 states. Flux is given in units of μs^−^1. The eleven states presented here correspond to: (1) GR24 productively bound (D-ring facing S97) and CTH in the associated state, (2) GR24 unproductively bound (ABC rings facing S97) and CTH in the associated state, (3) GR24 associated to T1/T4 and CTH in the associated state, (4) GR24 and CTH interaction with CTH in the associated state, (5) GR24 unbound and CTH in the associated state, (6) GR24 productively bound and CTH in the dissociated state, (7) GR24 unproductively bound and CTH in the dissociated state, (8) GR24 and CTH interaction in the T1/T4 associated state (CTH in its dissociated state), (9) GR24 associated with T1/T4 and CTH dissociated but not interacting with GR24, (10) GR24 and CTH non-T1/T4 associated state interaction with CTH in the dissociated state, (11) GR24 unbound and CTH in the dissociated state. The top two fluxes to each state is shown, with all of State 2 and fluxes to State 7 omitted due to their low relevance and to improve figure clarity. States where D14 is associated are bordered in black, while states where D14 is dissociated are bordered in red. A complete table of fluxes is presented in Table S9.

## Conclusions

Our large-scale MD simulations have demonstrated the interactions between D14 and D3 both with and without GR24 and shown the role these interactions play in inhibiting SL hydrolysis. We found that the CTH lowered the free-energy barrier for catalytic inactivation, which stabilized D14 in inactive conformations which cannot hydrolyze GR24. Therefore, D14-CTH associations allosterically inhibit D14 enzymatic activity. Additionally, we saw frequent contact between the CTH and GR24, especially in three-way interactions at the T1/T4 helices. While GR24-CTH interactions stabilized the CTH far from the associated state, thereby removing the allosteric inhibition of D14, GR24-CTH contact prevented the ligand from entering the binding pocket. Contact probability analysis revealed a high probability of GR24 interaction with the CTH, with contact between GR24 and at least 1 CTH residue occurring in over 24% of frames. In contrast, pocket binding occurred in only 3% of frames. GR24-CTH association combined with the lid helix interactions stabilized GR24 in positions away from the binding pocket. Finally, transition path theory analysis revealed high flux into GR24-CTH associated states, with much lower flux into the GR24 pocket-bound states.

The interactions between D14, the CTH, and GR24 agree well with the experimentally observed decrease in SL hydrolysis in the presence of the CTH. ^12^ The allosteric inhibition of D14 by the CTH stabilizes catalytically inactive conformations of the catalytic triad residues. While CTH dissociation increases in the presence of GR24, this increase results from strong interactions between the CTH and GR24 that prevent ligand entry into the binding pocket and subsequent hydrolysis. This is also corroborated by our TPT analysis, in which flux into GR24-CTH associated states was significantly higher than flux into pocket-bound states. Understanding the CTH’s effect on SL hydrolysis offers insights on designing SL-based mechanisms to improve host crop yield while limiting harmful effects of *Striga* germination.

## Methods

### Molecular Dynamics Simulations

#### Simulation Protocol

Both *apo* and *holo* systems were created from structure 6BRT in the Protein Data Bank. ^12^ The GR24 model was taken from its bound structure in *Os*D14 (PDB 5DJ5) and inserted at a random position in the holo system using Packmol. ^13,33^ Simulations were set up with AmberTools 18 and run with Amber 18 using the ff14SB force field.^34^ Water was described with the TIP3P model and GR24 with the generalized AMBER force field (GAFF). ^35,36^ Simulations were run in a 70 Å cube TIP3P water box at an NaCl concentration of 0.15 M to ensure neutral ionic charge. Structures were minimized with the conjugate gradient descent method for 10000 steps, then equilibrated for 5 ns. The thermostat kept simulations at a constant temperature of 300K and the barostat kept a constant pressure of 1.0 bar. Short-range non-bonded interactions were calculated with a cutoff of 10 Å and long-range electrostatics calculated with the Particle Mesh Ewald algorithm. ^37^ The SHAKE algorithm was used for constraining bonds to hydrogen. ^38^

#### Adaptive Sampling

Rather than run single long trajectories for each MD system, we used an adaptive sampling scheme in which each round of simulations was seeded from the least sampled conformations from the previous round of simulations. Sampling metrics and a summary of simulations run for each round are provided in Tables S1-S2 for the *apo* system and S3-S4 for the *holo* system.

### Trajectory Analysis

#### Feature Calculations

All contact information was calculated with the MDTraj analysis package version 1.9.4.^39^ Residue contact distances were calculated with residue alpha carbons, and GR24 features were calculated with specific atoms on the ligand. Center-of-mass distance was calculated from all the CTH residues. Helical content was calculated based on Kabsch-Sander secondary structure definitions.^40^

#### Contact Probability Calculation

Contact probability was calculated for both terminals of the CTH, as well as residue A279, chosen for observed high contact levels with A223 on D14. The probability of GR24 contacts with both D14 and the CTH were computed as well. This calculation shows what fraction of total frames has the residue of interest contacting each residue on D14 (plus each residue of the CTH for the GR24 calculation). Contact probabilities were weighted by MSM probabilities using Eq. 1.

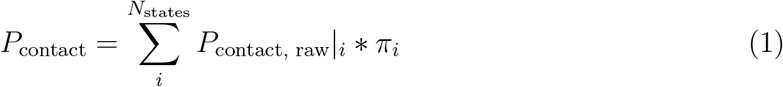

In addition to individual residue-ligand contact probabilities, overall probability of the GR24 ligand contacting the helix was calculated as

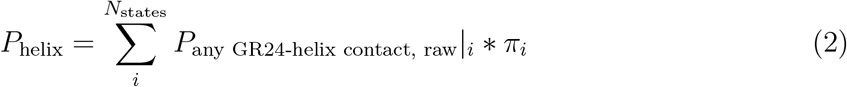

Overall probabilities of ligand binding were calculated using

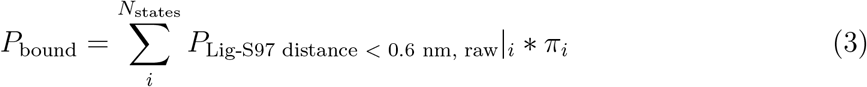

### Markov State Model Construction

Markov state models were built in pyEMMA from 21 total metrics involving D14-CTH interaction for both systems and GR24-protein interaction for the holo system. ^41^ A complete list of metrics is available in the Supporting Information. A hyperparameter search was performed using cross-validation to generate a VAMP1 score.^42^ Final MSM hyperparameters are listed in Table S5. Markovian behavior of MSMs was evaluated using the Chapman-Kolmogorov test (Fig. S6). Using the state equilibrium probabilities from Markov state models, free energy landscapes were calculated using

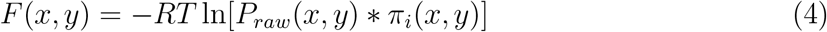

### Transition Path Theory

Twelve macrostates for the *holo* system were defined based on the relative positions of D14, the CTH, and GR24. Six positions were defined for GR24 and two for the CTH, and every combination of these two states were used. Definitions are given in Table S7. Based on the definitions, the three-way interaction between the D14 T1/T4 helices, the CTH, and GR24 was unpopulated, so the final calculation used only eleven states. These state definitions are given in Table S8. The mean first passage time (MFPT) was computed between each pair of macrostates.

## Supporting information

Supplementary Methods, Images, Tables and Results

## Supporting Information

This article contains supporting information. In-house code used for analysis is available at http://github.com/ShuklaGroup/Strigolactone-CTH-Binding.

## Acknowledgement

B.L.S. acknowledges support from the XSEDE scholars program and the Clare Boothe Luce scholars program. J.C. is a member of the NIH Chemistry-Biology Interface Training Program (T32-GM136629). D.S. acknowledges support from the CAS Fellowship, Center for Advanced Studies at University of Illinois at Urbana-Champaign and Sloan Research Fellowship from Alfred P. Sloan Foundation.

