## Supplementary Methods, Images, Tables and Results for "Dual role of strigolactone receptor signaling partner in inhibiting substrate hydrolysis"

### Details of adaptive sampling protocol

Table S1: Adaptive sampling metrics and selection criteria for *apo* system.

| Round | Sampling Metrics | Selection Criteria |
|---|---|---|
| 1 | - | Initial structure |
| 2 | A223-D3 N-terminus distance<br>A223-D3 C-terminus distance | 4 random clusters, 25 points per cluster |
| 3 | A223-D3 N-terminus distance<br>A223-C-terminus distance | 4 random clusters, 25 points per cluster |
| 4-9 | A223-D3 N-terminus distance<br>A223-D3 C-terminus distance<br>D3 orientation relative to D14 | 100 clusters,<br>5 points each from 20 least populated |
| 10-12 | A223-D3 N-terminus distance<br>A223-D3 C-terminus distance<br>D3 orientation relative to D14<br>Distance between CTH terminals | 100 clusters,<br>5 points each from 20 least populated |
| 13-14 | TICA coordinate 1<br>TICA coordinate 2 | 25 points each from 4 undersampled regions |
| 15 | TICA coordinate 3<br>TICA coordinate 5 | 25 points each from 4 undersampled regions |

Table S2: Summary of adaptive rounds for *apo* system

| Round | Parallel Trajectories | Trajectory length (ns) | Aggregate ($\mu$s) |
|---|---|---|---|
| 1 | 1 | 130 | 0.130 |
| 2 | 100 | 130 | 12.9 |
| 3 | 100 | 130 | 13.0 |
| 4 | 100 | 130 | 13.0 |
| 5 | 100 | 130 | 13.0 |
| 6 | 100 | 130 | 13.0 |
| 7 | 100 | 130 | 13.0 |
| 8 | 100 | 130 | 13.0 |
| 9 | 100 | 130 | 12.6 |
| 10 | 100 | 130 | 13.0 |
| 11 | 100 | 130 | 13.0 |
| 12 | 100 | 130 | 13.0 |
| 13 | 100 | 130 | 12.2 |
| 14 | 100 | 130 | 13.0 |
| 15 | 100 | 130 | 12.7 |
| | | Total ($\mu$s) | 180.5 |

Table S3: Adaptive sampling metrics and selection criteria for *holo* system.

| Round | Sampling Metrics | Selection Criteria |
|---|---|---|
| 1 | - | Initial structure |
| 2 | I120-D3 N-terminus distance<br>D-ring-S97 distance | 4 random clusters, 25 points per cluster |
| 3 | A223-D3 N-terminus distance<br>D-ring-S97 distance<br>A-ring-S97 distance | 100 clusters,<br>5 points each from 20 least populated<br>and D-ring-S97 distance < 3nm |
| 4 | A223-D3 N-terminus distance<br>D-ring-S97 distance<br>B-ring-L283 distance | 100 clusters,<br>5 points each from 20 least populated<br>and D-ring-S97 distance < 3nm |
| 5-6 | A223-D3 N-terminus distance<br>D-ring-S97 distance<br>B-ring-L283 distance<br>K230-A279 | 100 clusters,<br>5 points each from 20 least populated<br>and D-ring-S97 distance < 3nm |
| 7-8 | A223-D3 N-terminus distance<br>D-ring-S97 distance<br>B-ring-L283 distance<br>K230-A279 | 100 clusters,<br>5 points each from 20 least populated<br>and D-ring-S97 distance < 3nm |
| 9 | A223-D3 N-terminus distance<br>D-ring-S97 distance<br>B-ring-L283 distance<br>K230-A279<br>CTH helical content | 200 clusters,<br>5 points each from 40 least populated |
| 10 | TICA coordinate 1<br>TICA coordinate 3 | 50 points each from 4 undersampled regions |
| 11 | S220-H247 distance<br>TICA coordinate 4 | 200 points from undersampled region |
| 12 | TICA coordinate 1<br>TICA coordinate 3 | 200 points from undersampled region |

Table S4: Summary of adaptive rounds for *holo* system

| Round | Parallel Trajectories | Trajectory length (ns) | Aggregate ($\mu$s) |
|---|---|---|---|
| 1 | 1 | 120 | 0.120 |
| 2 | 100 | 120 | 12.0 |
| 3 | 100 | 120 | 12.0 |
| 4 | 100 | 120 | 12.0 |
| 5 | 100 | 120 | 12.0 |
| 6 | 100 | 120 | 12.0 |
| 7 | 100 | 120 | 12.0 |
| 8 | 200 | 120 | 12.0 |
| 9 | 200 | 120 | 24.0 |
| 10 | 200 | 120 | 23.6 |
| 11 | 200 | 120 | 24.0 |
| 12 | 200 | 120 | 23.3 |
| 13 | 200 | 120 | 23.3 |
| | | Total ($\mu$s) | 202.3 |

### Markov state model construction and validation

#### MSM Features

To construct our Markov state models, we first calculated a set of inter-residue distance and helical content features from our simulation data. The full set of features used for MSM construction is shown in Table S5.

Table S5: Featurizations used for MSM construction. C-$\alpha$ distances were used for all inter-residue distances.

| | *Apo* | *Holo* |
|---|---|---|
| CTH Dissociation | A223-D3 N-terminus<br>A223-D3 C-terminus<br>K230-A279<br>A223-E276<br>L283-A223<br>R284-K230 | A223-D3 N-terminus<br>A223-D3 C-terminus<br>K230-A279 |
| CTH-T1 distance | G149-R284 | - |
| CTH-T4 distance | F195-D3 N Terminus | - |
| CTH-D14 Contacts | R267-D3 N-terminus<br>T238-D3 C-terminus<br>Y132-T275<br>E240-L283 | D131-D3 C-terminus |
| CTH Structure | Helical Content<br>N-C Terminal Distance | Helical Content<br>N-C Terminal Distances |
| D-loop contacts | T216-H247<br>R217-H247<br>D218-H247<br>V219-H247<br>S220-H247<br>V221-H247<br>P222-H247 | T216-H247<br>R216-H247<br>D217-H247<br>I218-H247<br>M219-H247<br>V220-H247<br>P221-H247 |
| Ligand - catalytic triad | -<br>- | A-ring-S97<br>D-ring-S97 |
| Ligand - D14 | -<br>-<br>- | A-ring-A223<br>D-ring-A223<br>D-ring-F195 |
| Ligand - CTH | -<br>- | D-ring-G285<br>B-ring-L283 |

#### MSM hyperparameter selection

To select lag times for our MSMs, we plotted implied timescales for models estimated at a series of lag times and chose a lag time at which the implied timescale stopped changing with increasing lag time. Implied timescale plots are shown in Fig. S1.

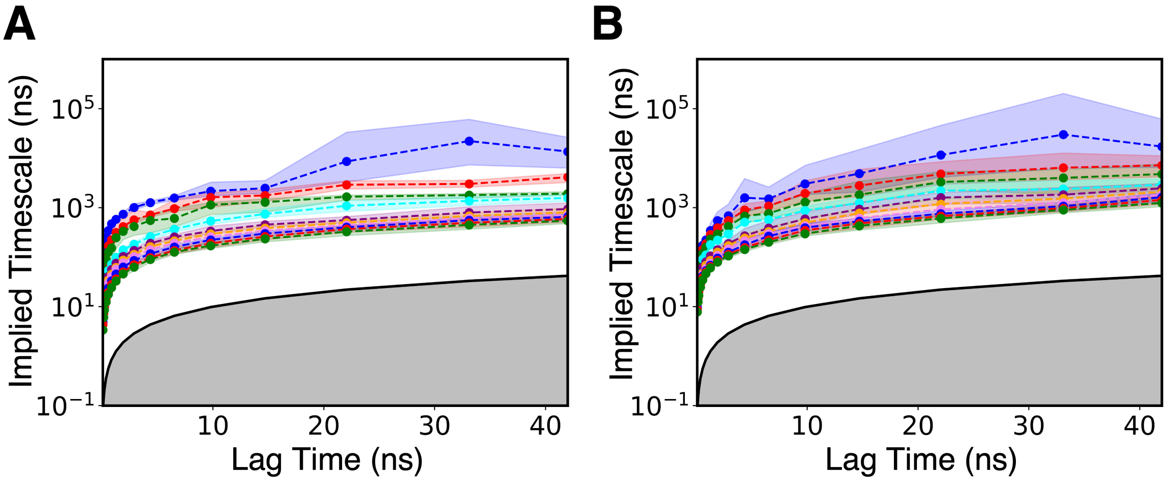


Figure S1: Implied timescale plots for the *apo* (A) and *holo* (B) systems. Lag time was chosen as 15 ns for both systems.

To select number of TICA components and number of clusters to discretize our simulation data for MSM construction, we performed a grid search in which we calculated a cross-validation score for MSMs calculated with different parameter sets. Cross-validation scores for different parameter sets are shown in Fig. S2. Final parameters used for MSM construction are shown in Table S6.

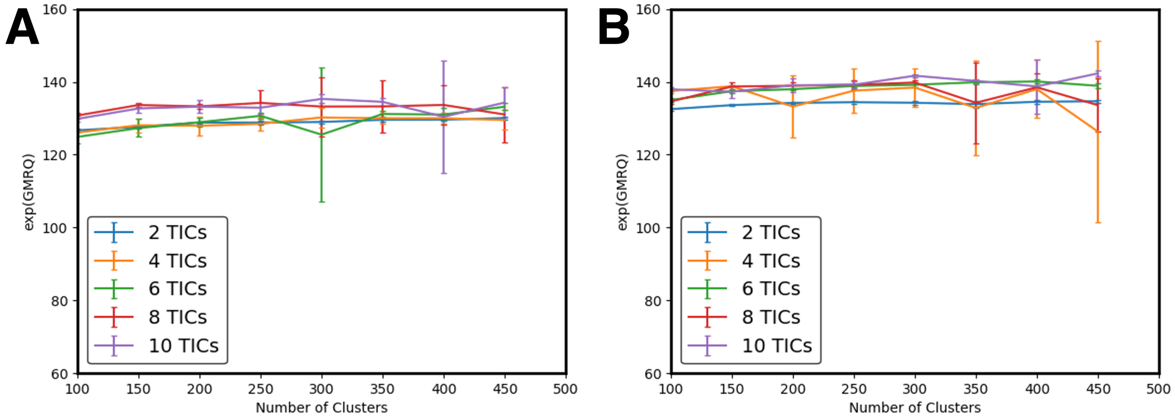


Figure S2: Cross-validation scores calculated with different hyperparameter sets for the *apo* (A) and *holo* (B) systems.

Table S6: Final parameters used for MSM construction

| | *Apo* | *Holo* |
|---|---|---|
| Lag time (ns) | 15 | 15 |
| Number of TICA components | 10 | 6 |
| Number of clusters | 350 | 400 |

#### Chapman-Kolmogorov validation

To validate our MSMs, we employed the Chapman-Kolmogorov test. Briefly, if a system exhibits Markovian behavior, the $n$th power of a transition matrix estimated at lag time $\tau$ should equal the transition matrix estimated at lag time $n\tau$. Results of the Chapman-Kolmogorov test are shown in Fig. S3.

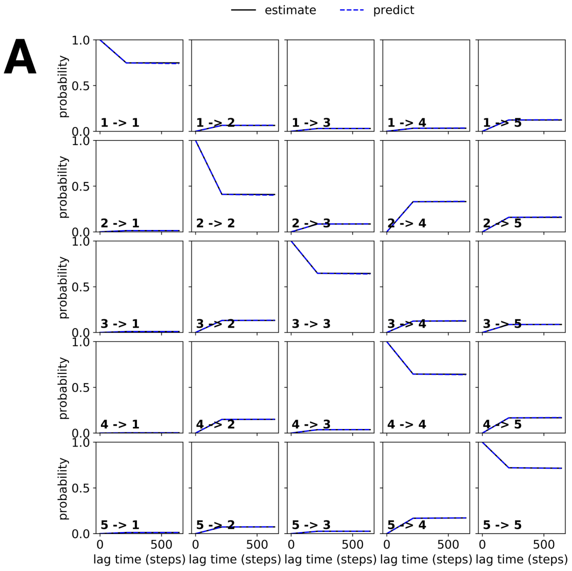


Figure S3: Chapman-Kolmogorov tests for the *apo* (A) and *holo* (B) systems. The test shows good agreement between estimated and predicted MSMs, indicating that the models follow the Markov property.

### Transition path theory

During transition path theory calculation for the *holo*, we coarse-grained our data by assigning MSM clusters to a set of eleven macrostates defined by positions of the GR24 ligand and CTH. Definitions for different ligand and CTH positions are shown in Table S7 and macrostates are shown in Table S8. All calculated fluxes between the eleven macrostates are shown in Table S9.

Table S7: Definitions of ligand binding and CTH association used for TPT calculation

| State | Parameters |
|---|---|
| GR24 Bound | Minimum D-ring-S97 distance $<$ 0.6 nm<br>AND<br>Mean A-ring-S97 distance $>$ Mean D-ring-S97 distance<br>AND<br>Not included in T1/T4 associated state |
| GR24 Inversely Bound | Mean A-ring-S97 distance $<$ 1.5 nm<br>AND<br>Mean D-ring-S97 distance $<$ 1.5 nm<br>AND<br>Not included in T1/T4 associated state |
| GR24-T1/T4 Association | Minimum A-ring-E138 distance $<$ 1 nm<br>AND<br>D-ring-F196 distance $<$ 1.3 nm |
| GR24-CTH Association | Minimum B-ring-A279 distance $<$ 0.9 nm<br>AND<br>B-ring-L283 distance $<$ 0.9 nm |
| GR24 Unbound | Not included in any other GR24 state |
| CTH-D14 Association | Minimum N-term-A223 distance $<$ 2.0 nm<br>AND<br>Minimum C-Term-K230 distance $<$ 1.4 nm<br>AND<br>Minimum C-term-V239 distance $<$ 1.9 nm<br>AND<br>Minimum A279-A223 distance $<$ 1.2 nm |
| CTH-D14 Dissociation | Not included in CTH-D14 associated state |

Table S8: Definitions of the eleven macrostates used for TPT calculation

| State Number | Definition |
|---|---|
| 1 | GR24 Bound<br>CTH Associated |
| 2 | GR24 Inverse<br>CTH Associated |
| 3 | GR24+T1/T4<br>CTH Associated |
| 4 | GR24+CTH<br>CTH Associated |
| 5 | GR24 Unbound<br>CTH Associated |
| 6 | GR24 Bound<br>CTH Dissociated |
| 7 | GR24 Inverse<br>CTH Dissociated |
| 8 | GR24+T1/T4+CTH<br>CTH Dissociated |
| 9 | GR24+T1/T4<br>CTH Dissociated |
| 10 | GR24+CTH<br>CTH Dissociated |
| 11 | GR24 Unbound<br>CTH Dissociated |

Table S9: Rate constants of flux between TPT states (µs$^-1$)

| | 1 | 2 | 3 | 4 | 5 | 6 | 7 | 8 | 9 | 10 | 11 |
|---|---|---|---|---|---|---|---|---|---|---|---|
| 1 | - | 0.27 | 1.25 | 0.82 | 5.61 | 0.18 | 0.26 | 0.36 | 0.80 | 1.30 | 2.30 |
| 2 | 0.32 | - | 1.02 | 0.83 | 1.45 | 0.18 | 0.26 | 0.42 | 0.63 | 0.90 | 1.65 |
| 3 | 0.43 | 0.31 | - | 1.58 | 6.89 | 0.22 | 0.35 | 0.57 | 1.49 | 2.68 | 5.52 |
| 4 | 0.41 | 0.32 | 3.02 | - | 5.61 | 0.20 | 0.33 | 0.56 | 1.48 | 2.42 | 6.32 |
| 5 | 1.06 | 0.33 | 2.94 | 1.62 | - | 0.22 | 0.35 | 0.64 | 1.49 | 2.34 | 4.98 |
| 6 | 0.42 | 0.28 | 4.42 | 1.45 | 7.52 | - | 0.32 | 0.51 | 1.28 | 2.96 | 10.24 |
| 7 | 0.42 | 0.30 | 4.56 | 1.72 | 8.42 | 0.21 | - | 0.54 | 1.26 | 3.12 | 6.58 |
| 8 | 0.39 | 0.32 | 2.99 | 1.61 | 6.85 | 0.21 | 0.33 | - | 1.23 | 2.32 | 6.02 |
| 9 | 0.43 | 0.28 | 2.84 | 1.45 | 6.14 | 0.19 | 0.29 | 0.48 | - | 2.43 | 5.16 |
| 10 | 0.45 | 0.29 | 2.74 | 1.50 | 6.39 | 0.21 | 0.33 | 0.52 | 1.30 | - | 4.94 |
| 11 | 0.42 | 0.36 | 2.93 | 1.73 | 5.20 | 0.22 | 0.33 | 0.59 | 1.42 | 2.38 | - |
